# Removal of Scanner Effects in Covariance Improves Multivariate Pattern Analysis in Neuroimaging Data

**DOI:** 10.1101/858415

**Authors:** Andrew A. Chen, Joanne C. Beer, Nicholas J. Tustison, Philip A. Cook, Russell T. Shinohara, Haochang Shou, the Alzheimer’s Disease Neuroimaging Initiative

## Abstract

To acquire larger samples for answering complex questions in neuroscience, researchers have increasingly turned to multi-site neuroimaging studies. However, these studies are hindered by differences in images acquired across multiple scanners. These effects have been shown to bias comparison between scanners, mask biologically meaningful associations, and even introduce spurious associations. To address this, the field has focused on harmonizing data by removing scanner-related effects in the mean and variance of measurements. Contemporaneously with the increase in popularity of multi-center imaging, the use of multivariate pattern analysis (MVPA) has also become commonplace. These approaches have been shown to provide improved sensitivity, specificity, and power due to their modeling the joint relationship across measurements in the brain. In this work, we demonstrate that methods for removing scanner effects in mean and variance may not be sufficient for MVPA. This stems from the fact that such methods fail to address how correlations between measurements can vary across scanners. Data from the Alzheimer’s Disease Neuroimaging Initiative is used to show that considerable differences in covariance exist across scanners and that popular harmonization techniques do not address this issue. We also propose a novel methodology that harmonizes covariance of multivariate image measurements across scanners and demonstrate its improved performance in data harmonization.

## 1 Introduction

The need for larger samples in human subjects research have led to a growing number of multi-site studies that aggregate data across multiple locations. This trend is especially prevalent in neuroimaging research where the reliability and generalizabilty of findings from the conventional single-site studies are often limited by the ability to recruit and study sufficiently large and representative samples from the population. Many consortia have been formed to address such issues (Mueller et al., 2005; Sudlow et al., 2015; Trivedi et al., 2016; Van Essen et al., 2013). The larger samples obtained through these efforts promote greater power to detect significant associations as well as better generalizability of results. However, these study designs also introduce heterogeneity in acquisition and processing that, if not appropriately addressed, may impact study findings.

Several researchers have determined that variability driven by scanner, often called scanner effects, reduce the reliability of derived measurements and can introduce bias. Neuroimaging measurements have been repeatedly shown to be affected by scanner manufacturer, model, magnetic field strength, head coil, voxel size, and acquisition parameters (Han et al., 2006; Kruggel et al., 2010; Reig et al., 2009; Wonderlick et al., 2009). Yet even in scanners of the exact same model and manufacturer, differences still exist for certain neuroimaging biomarkers (Takao et al., 2011).

Until recently, neuroimaging analyses primarily involved mass univariate testing which treats features as independent and does not leverage covariance between features. Under this paradigm, the impact of scanner effects is through changes in the mean and variance of measurements. Increasingly, researchers have used sets of features as patterns for prediction algorithms or traditional multivariate methods in a framework called multivariate pattern analysis (MVPA). This approach has become a powerful tool in diverse research topics including pain perception (Smith et al., 2017), neural representations (Haxby et al., 2014), and psychiatric illnesses (Koutsouleris et al., 2014). One of the major benefits of MVPA is that it leverages the joint distribution and correlation structure among multivariate brain features in order to better characterize a phenotype of interest (O’Toole et al., 2007). As a result, scanner effects on the covariance of measurements are likely to impact findings substantially. In fact, a recent investigation showed that MVPA was able to detect scanner with high accuracy and that the detection of sex depended heavily on the scanners included in the training and test data (Glocker et al., 2019).

The major statistical harmonization techniques employed in neuroimaging have generally corrected for differences across scanners in mean and variance, but not covariance (Fortin et al., 2016, 2018; Rao et al., 2017; Yamashita et al., 2019). Increasingly, the ComBat model (Johnson et al., 2007) has become a popular harmonization technique in neuroimaging and has been successfully applied to structural and functional measures (Bartlett et al., 2018; Fortin et al., 2017, 2018; Marek et al., 2019; Yu et al., 2018). However, this model does not address potential scanner effects in covariance.

Recently, another stream of data-driven harmonization methods have aimed to apply generative adversarial networks (GANs) or distance-based methods to unify distributions of measurements across scanners. However, the GAN-based harmonization methods have mostly been tested for image-level harmonization only and are limited in options to protect for clinical effects in the data (Gao et al., 2019; Nguyen et al., 2018; Zhong et al., 2020). A recent distance-based method is applicable to derived measurements and has been tested in classification of Alzheimer’s disease using support vector machines (Zhou et al., 2018). However, the method in Zhou et al. (2018) has not been tested for detection of site via MVPA and also requires several conditions which may not hold in studies with sufficiently heterogenous scanners or major differences in subject demographics across scanners.

In this paper, we examine whether scanner effects influence MVPA results. In particular, we study the cortical thickness measurements derived from images acquired by the Alzheimer’s Disease Neuroimaging Initiative (ADNI) and demonstrate the existence of scanner effects in covariance of structural imaging measures. We then propose a novel harmonization method called Correcting Covariance Batch Effects (CovBat) that removes scanner effects in mean, variance, and covariance. We apply CovBat and show that within-scanner correlation matrices are successfully harmonized. Furthermore, we find that machine learning methods are unable to distinguish scanner manufacturer after our proposed harmonization is applied, and that the CovBat-harmonized data facilitate more accurate prediction of disease group. We also assess the performance of the proposed method in simulated data, and again find that the method mitigates scanner effects and improves detection of meaningful associations. Our results demonstrate the need to consider covariance in harmonization methods, and suggest a novel procedure that can be applied to better harmonize data from multi-site imaging studies.

## 2 Methods

### 2.1 ADNI data analysis

All data for this paper are obtained from ADNI (http://adni.loni.usc.edu/ and processed using the ANTs longitudinal single-subject template pipeline (Tustison et al., 2019) with code available on GitHub (https://github.com/ntustison/CrossLong). Informed consent was obtained by all participants in the ADNI study. Institutional review boards approved the study at all of the contributing institutions.

We briefly summarize the steps involved. First, we download raw T1-weighted images from the ADNI-1 database, which were acquired using MPRAGE for Siemens and Philips scanners and a works-in-progress version of MPRAGE on GE scanners (Jack et al., 2010). For each subject, we estimate a single-subject template from all the image timepoints. After rigid spatial normalization to this single-subject template, each normalized timepoint image is then processed using the single image cortical thickness pipeline consisting of brain extraction (Avants et al., 2010), denoising (Manjón et al., 2010), N4 bias correction (Tustison et al., 2010), Atropos n-tissue segmentation (Avants et al., 2011), and registration-based cortical thickness estimation (Das et al., 2009). For our analyses, we use the 62 cortical thickness values from the baseline scans. The covariance matrices for these cortical thicknesses in the three largest sites are shown in Fig. 1.

**Figure 1:**
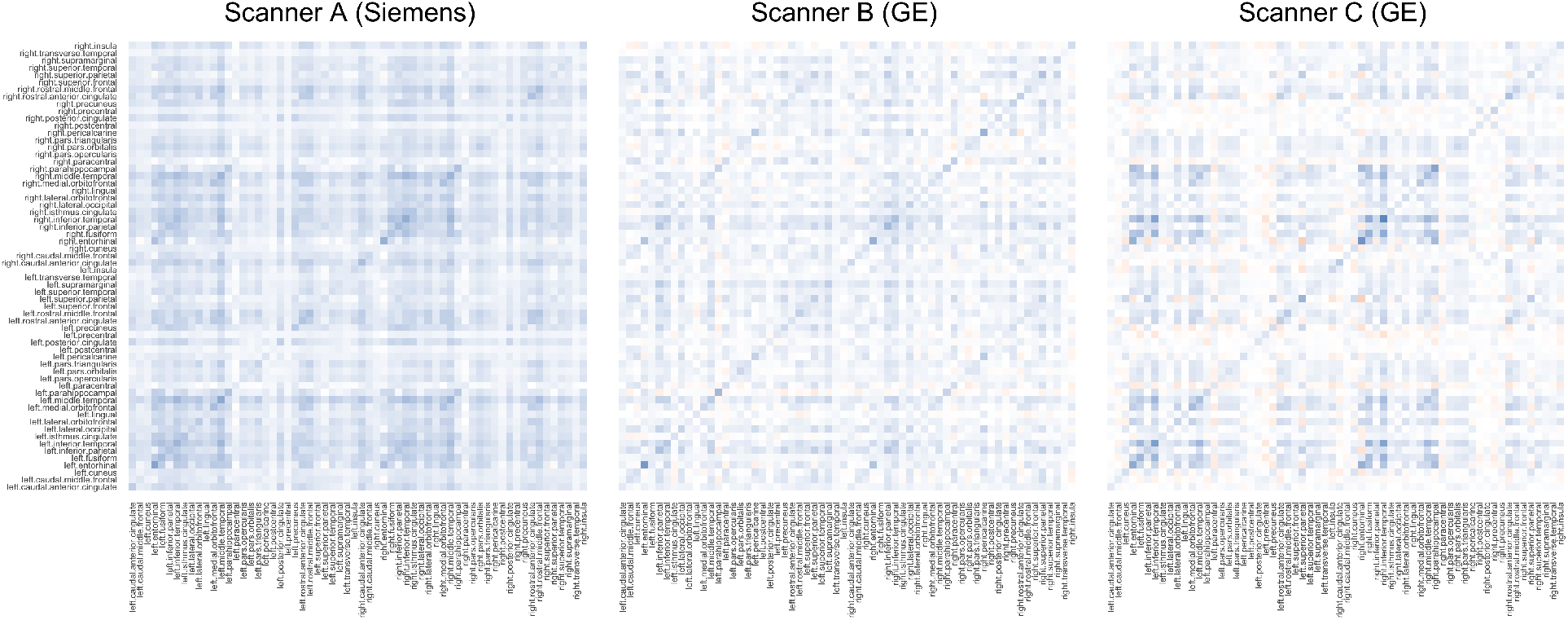
Covariance matrices for cortical thickness values acquired on three scanners. Scanner A was a Siemens Symphony 1.5T scanner and the other scanners are General Electric Signa Excite 1.5T scanners.

We define scanner based on information contained within the Digital Imaging and Communications in Medicine (DICOM) headers for each scan. Specifically, subjects are considered to be acquired on the same scanner if they share the scanner site, scanner manufacturer, scanner model, head coil, and magnetic field strength. In total, this definition yields 142 distinct scanners of which 78 had less than three subjects and were removed from analyses. The final sample consists of 505 subjects across 64 scanners, with 213 subjects imaged on scanners manufactured by Siemens, 70 by Philips, and 222 by GE. The sample has a mean age of 75.3 (SD 6.70) and is comprised of 278 (55%) males, 115 (22.8%) Alzheimer’s disease (AD) patients, 239 (47.3%) late mild cognitive impairment (LMCI), and 151 (29.9%) cognitively normal (CN) individuals.

The ADNI sample demographics are considerably different across sites, which precludes application of certain harmonization methods. For example, Zhou et al. (2018) relies on ‘‘nontrivial overlap” of the potential confounders across sites and proposed a subsampling approach that performs distributional shifts on subsamples of data matched by the discrete stratum of the confounders. Given that our data are sufficiently heterogeneous, it is challenging to form bins matched by age, sex, and diagnosis status to ensure that each site has at least one individual in each bin. This prevents protection of age effects in applying the harmonization method proposed by Zhou et al. (2018).

### 2.2 Combatting Batch Effects

We first review ComBat (Fortin et al., 2017, 2018; Johnson et al., 2007) for harmonization of neuroimaging measures. ComBat seeks to remove the mean and variance scanner effects of the data in an empirical Bayes framework. Let ***y**_ij_* = (*y*_*ij*1_, *y*_*ij*2_,…, *y_ijp_*)^*T*^, *i* = 1, 2,…, *M, j* = 1, 2,…, *n_i_* denote the *p* × 1 vectors of observed data where *i* indexes scanner, *j* indexes subjects within scanners, *n_i_* is the number of subjects acquired on scanner *i*, and *p* is the number of features. Our goal is to harmonize these vectors across the *M* scanners. ComBat assumes that the features indexed by *v* follow

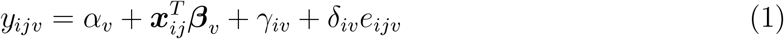

where *α_v_* is the intercept, ***x**_ij_* is the vector of covariates, ***β**_v_* is the vector of regression coefficients, *γ_iv_* is the mean scanner effect, and *δ_iv_* is the variance scanner effect. The errors *e_ijv_* are assumed to follow 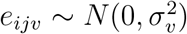. ComBat finds least-squares estimates 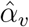 and 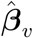 for each feature and empirical Bayes point estimates 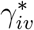 and 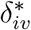 for each feature across sites by pooling information across features. Finally, ComBat residualizes with respect to these estimates to obtain harmonized data

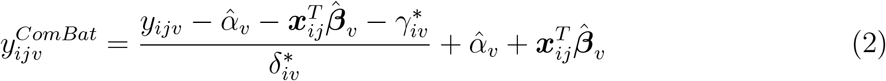

### 2.3 Correcting Covariance Batch Effects

To address potential covariance scanner effects, we build on the existing ComBat framework. We again assume that the features follow

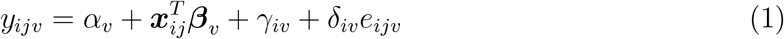

However, the error vectors ***e**_ij_* = (*e*_*ij*1_, *e*_*ij*2_,…, *e_ijp_*)^*T*^ ~ *N*(**0**, Σ_*ij*_) may be spatially correlated and differ in covariance across scanner. Analogous to how ComBat modifies observations to bring each within-scanner variance to the pooled variance across sites, our proposed method modifies principal component (PC) scores to shift each within-scanner covariance to the pooled covariance structure. We achieve this by approximating within-scanner covariance structures using the principal components and PC scores obtained across all observations. We propose the CovBat algorithm, which accounts for the joint distribution of ComBat-adjusted observations as follows:

**Step 1.** We first perform ComBat to remove the mean and variance shifts in the marginal distributions of the cortical thickness measures. Then, we additionally residualize with respect to the intercept and covariates to obtain ComBat-adjusted residuals denoted 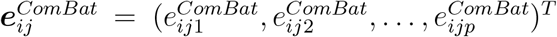 where *p* is the number of features. We then define these residuals using notation from **Section 2.2** as

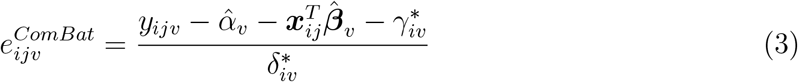

where *i* = 1, 2,…, *M, j* = 1, 2,…, *n_i_, M* is the number of scanners, and *n_i_* is the number of subjects acquired on scanner *i*. 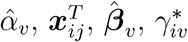, and 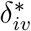 are defined in **Section 2.2**.
**Step 2.** The 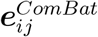 are assumed to have mean 0; their covariance matrices which we denote by Σ_*i*_, however, may differ across scanners. We first perform principal components analysis (PCA) on the full data residuals and represent the full data covariance matrix as 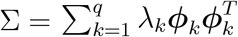 where the rank 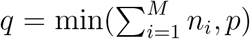, λ_*k*_ are the eigenvalues of Σ, and *ϕ_k_* are the principal components obtained as the eigenvectors of Σ. In practice, PCA is performed on the sample covariance matrix 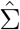 and we obtain estimated eigenvalues 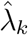 and eigenvectors 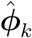. The ComBat-adjusted residuals can then be expressed as 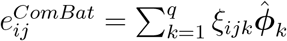 where *ξ_ijk_* are the principal component scores. We then aim to bring each within-scanner covariance matrix Σ_*i*_ to the pooled covariance across scanners. Since our goal is to recover covariance structures resembling Σ, we approximate the within-scanner covariance matrices as 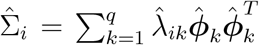 where 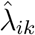 are within-scanner eigenvalues estimated as the sample variance of the principal component scores 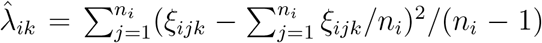 and 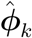 are estimated from the full data covariance. This model assumes that the covariance scanner effect is contained within the variances of the PC scores with the principal components estimated from the full data.
**Step 3.** Thus, we posit:

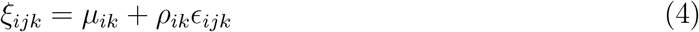

where 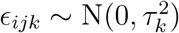, *τ_k_* is the variance of the *k*th score and *μ_ik_, ρ_ik_* are the center and scale parameters corresponding to principal components *k* = 1, 2,… *K* where *K* ≤ *q* is a tuning parameter chosen to capture the desired proportion of the variation in the observations. If *K* is chosen such that *K* = *q*, all principal components are harmonized. Note that this is analogous to the ComBat model, applied to each of the *k* principal component scores instead of the original measures. We can then estimate each of the *K* pairs of center and scale parameters by finding the values that bring each scanner’s mean and variance in scores to the pooled mean and variance, which we denote 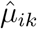) and 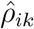. We then remove the scanner effect in the scores via 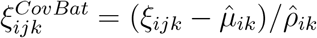.
**Step 4.** We obtain CovBat-adjusted residuals 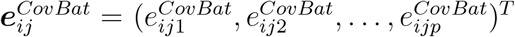 by projecting the adjusted scores back into the residual space via

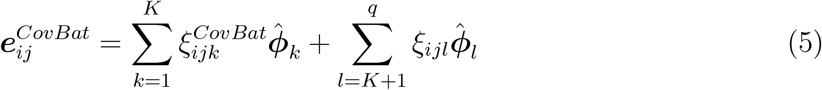

We then add the intercepts and covariates effects estimated in Step 1 to obtain CovBat-adjusted observations

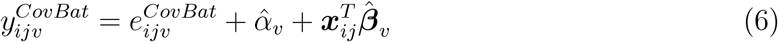

### 2.4 Multivariate pattern analysis experiments

Using the full ADNI sample, we evaluate whether the harmonization procedures affect the results of MVPA using the neuroimaging measures as patterns for a prediction algorithm. We achieve this through a Monte Carlo split-sample experiment where we i) randomly split the subjects into 50% training set and 50% validation set, ii) train a random forests algorithm to detect either scanner manufacturer or a binary clinical covariate, and iii) assess predictive performance on the validation set via AUC. We train separate models for unharmonized, ComBat-harmonized, and CovBat-harmonized data where both harmonization methods are performed including age, sex, and diagnosis status as covariates. CovBat harmonization is performed using 37 PCs, which cumulatively explain 95% of the variation. We perform steps (i)-(iii) 100 times for each dataset and report the mean AUC along with standard deviation. For these experiments, lower AUC for detection of scanner manufacturer and higher AUC for detection of clinical covariates would indicate improved harmonization. For prediction of scanner manufacturer, we avoid the possibility that scanner could be detected through the covariates age, sex, and disease status by residualizing out these variables from each cortical thickness value via linear models.

We also evaluate the harmonization methods using multivariate analysis of variance (MANOVA), a more classical form of MVPA. MANOVA tests for differences in mean across groups in multivariate data, but is known to be sensitive to differences in covariance across groups. We perform MANOVA across scanner manufacturer, sex, and diagnosis status using Pillai’s trace, which is known to be more robust to inhomogeneity in covariance than other alternative test statistics (Olson, 1974). We report *p*-values for these associations before and after harmonization.

### 2.5 Simulation design

Let ***y**_ij_, i* = 1,2, 3, *j* = 1,2,…, *n_i_* be vectors of length *p* representing simulated cortical thickness values for three sites, each with *n_i_* observations. The ***y**_ij_* are generated using the following model:

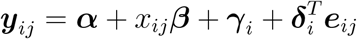

where ***α*** is the first *p*/2 elements in each hemisphere from the sample mean vector of Scanner B observations in the ADNI data, ***β*** is the vector of predictor effects on the mean, *x_ij_* is a binary covariate drawn from a Bernoulli distribution with probability 0.25, and ***e**_ij_* is the vector of error terms. The mean site effects *γ_i_* = (*γ*_*i*1_, *γ*_*i*2_,…, *γ_ip_*)^*T*^ are vectors drawn from independent and identically distributed (i.i.d.) normal distributions with mean zero and standard deviation 0.1. The variance site effects ***δ**_i_* = (*δ*_*i*1_, *δ*_*i*2_,…, *δ_ip_*)^*T*^ are vectors drawn from site-specific inverse gamma distributions with chosen parameters. For our simulations, we chose to distinguish the site-specific scaling factors by assuming 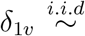 Inverse Gamma (46, 50), 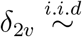 Inverse Gamma (51, 50), and 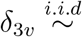 Inverse Gamma (56, 50) for *v* = 1, 2,…, *p*.

#### 2.5.1 Simple covariance effects

We first assess whether CovBat can recover the underlying covariance structure when the covariance site effects are captured by its PC directions. We refer to this simulation setting as the Simple Covariance Effects simulation. For *p* = 62, we set the underlying covariance *S* as the sample correlation matrix of cortical thickness observations in the ADNI data, with eigendecomposition 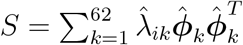. We then generate error terms ***e**_ij_* that contain site-specific shifts in the first eigenvalue via

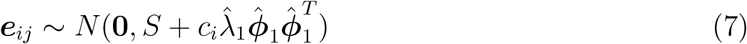

where *c_i_* controls the severity of the covariance shift. For our simulations, we set *c*_1_ = −1/2, *c*_2_ = 0, and *c*_3_ = 1/2 so that the pooled covariance structure is equivalent to *S*. We choose *β_i_* = −0.5 for ⌊−*p*/4⌋ regions of interest in both the left and right hemispheres to associate the predictor with decreases in mean simulated cortical thicknesses.

For several choices of within-site sample size and number of features, we generate 1000 datasets using the above model and evaluate recovery of the underlying covariance structure across sites and harmonization methods. For each site, we calculate the average Frobenius distance across datasets between each sample within-site covariance matrix and the true covariance S. We then report the average across sites before and after harmonization, where CovBat harmonization is performed on PCs that explain 95% of the variation. We also perform MANOVA to test associations with site and the simulated predictor to evaluate if the harmonization methods can successfully remove site effects while preserving associations of interest. For MANOVA, we report the rejection rate at a type I error rate of 0.05 across the 1000 datasets.

#### 2.5.2 Complex covariance effects

To evaluate how CovBat performs when the covariance site effects are not easily captured by the principal components and when the predictor may affect covariance, we modify the simulation to incorporate high-rank covariance shifts due to site via Ω_*i*_ and due to predictor as Ψ. For *p* = 62, the error terms ***e**_ij_* are now generated via

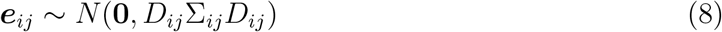

where Σ_*ij*_ = *S*+*x_ij_*Ψ+Ω_*i*_, *S* is the sample correlation matrix of cortical thickness observations in the ADNI data, Ψ is a chosen predictor-driven covariance shift matrix, and Ω_*i*_ are site-specific covariance shift matrices. The matrices *D_ij_* = **diag**(***d**_ij_*) where 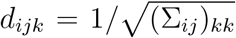 for *k* = 1, 2,…, 62 ensure that these covariance effects do not modify the marginal variances of ***e**_ij_*. To constrain the covariance matrices to be positive definite, we set the negative eigenvalues of Σ_*ij*_ equal to a small constant, 10^-12^. For *p* < 62, we instead generate ***e**_ij_* from the *p* × *p* submatrices of *D_ij_*Σ_*ij*_*D_ij_* constructed from the rows and columns corresponding to the first *p*/2 features in each hemisphere.

In this simulation scenario, *S* is no longer the pooled covariance structure since the covariance site effects Ω_*i*_ can take any form and these site-specific covariance structures do not necessarily combine across sites to resemble *S*. Instead of focusing on recovery of an underlying structure, we evaluate scanner effects throughout these simulations via MVPA. For several choices of within-site sample size and number of features, we generate 1000 datasets using the above model and perform experiments to test for scanner and covariate associations under a similar paradigm as the MVPA experiments implemented on the ADNI dataset. For each dataset, we i) randomly split the sample into 50% training data and 50% testing data, ii) train a random forests algorithm to recognize if the observations either are from Site 1 or have the simulated predictor, and iii) assess predictive performance on the testing data. Lower AUC for detection of Site 1 and higher AUC for detection of the simulated predictor indicate improved harmonization. CovBat harmonization is performed on PCs that explain 95% of the variation. For prediction of Site 1, we avoid the possibility that site could be detected through the predictor by using linear models to residualize out the predictor from each simulated cortical thickness value. For simulated datasets where the training set does not contain observations with the simulated predictor, the random forest algorithm cannot be trained so we generate another dataset to replace it. We also perform MANOVA for testing associations with site and simulated predictor and report the rejection rate at a type I error rate of 0.05 across the 1000 datasets.

We then design four simulation experiments to test how CovBat performs in multiple settings with varied covariance effects. In our ComBat simulation, we generate data without any covariance site effect from a model that resembles the ComBat model. In the Predictor Affects Mean simulation, we then introduce a covariance site effect to assess if CovBat can outperform ComBat in harmonization of covariance. We next introduce a predictor effect on covariance in the Predictor Affects Covariance simulation, which better illustrates how the detection of the predictor is affected by related covariance effects. Finally, we design a Covariance Only simulation which has no site or predictor effects in mean or variance, but still contains site and predictor effects in covariance. This final simulation illustrates how effects on covariance can influence MVPA results and be addressed through our proposed harmonization method.”

In the ComBat simulation, we impose site effects in mean and variance while having the predictor affect only the mean of the observations. We choose *β_i_* = −0.5 for ⌊*p*/4⌋ regions of interest in both the left and right hemispheres to impose that about half of the ROIs are negatively associated with the predictor. We also choose the Ω_*i*_ and Ψ to be 62 × 62 zero matrices to ensure that the covariance does not depend on site or the predictor.

In the Predictor Affects Mean simulation, we again impose that the predictor only affects the mean of the measurements but also introduce a site effect in covariance. We keep the same *β* and Ψ as in the ComBat simulation. However, we choose Ω_*i*_ to be distinct 62 × 62 high-rank matrices to distinguish covariance structures across sites.

In the Predictor Affects Covariance simulation, we assume that the predictor affects not only mean, but also covariance. We choose the predictor effect on covariance to be proportional to a site’s covariance shift. This scenario represents a situation where detection of the predictor using MVPA could be highly influenced by the presence of site effects. We use the same *β* value as in the ComBat and Predictor Affects Mean simulations but choose Ψ to be related to Ω_3_ to force confounding of Site 3 and predictor effects on covariance. To achieve this, we set Ψ = −(3/4)Ω_3_.

In the Covariance Only simulation, we assume that both site and the predictor influence the covariance, not the mean or variance. We fix *γ* = 0 and *δ* = **1** to remove site effects in mean and variance while also using the same Ω_*i*_ as in the Predictor Affects Mean and Predictor Affects Covariance simulations. Furthermore, we modify the predictor effect by setting *β* = **0** while keeping Ψ = –(3/4)Ω_3_ for the predictor effect in covariance.

## 3 Results

### 3.1 CovBat reduces covariance scanner effect

We first apply CovBat to observations acquired on the three scanners with the largest number of subjects. Scanner A consists of 23 subjects acquired on a Siemens Symphony 1.5T scanner while scanners B and C each consist of 20 subjects acquired on GE Signa Excite 1.5T scanners. See Table 1 for demographic details. CovBat is performed correcting 23 PCs, which explain 95% of the variation. For comparison, we also perform ComBat on the observations to obtain three datasets: unharmonized, ComBat-adjusted, and CovBat-adjusted. To ensure fair comparison of the covariance matrices across scanners, we residualize the cortical thickness measures across the three scanners jointly on age, sex, and diagnosis status while controlling for scanner in each dataset using a linear model. Fig. 2 shows the covariance matrices for each scanner using the residualized cortical measures both before and after harmonization. The differences between the unharmonized covariance matrices are striking. Especially notable are the increased positive covariance across most pairs of cortical regions in Scanner A and the weakened correlation between the right and left hemispheres in Scanner C, visible as the diagonal line in the top-left and bottom-right quadrants. Visually, the covariance differences remain similar after applying ComBat. These inter-scanner differences are considerably mitigated after CovBat.

**Figure 2:**
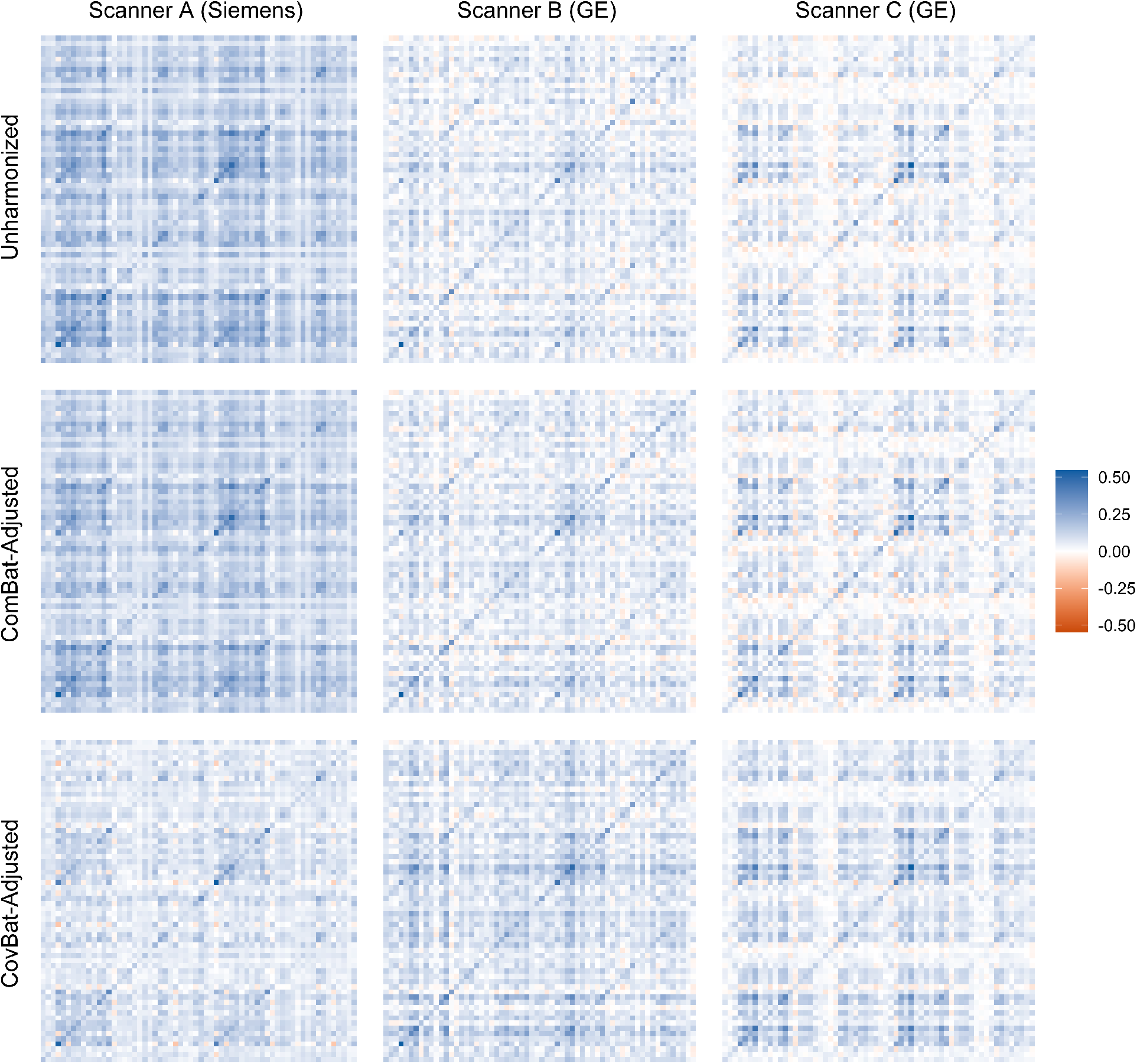
Covariance matrices for cortical thickness values acquired on three scanners before and after harmonization. All covariance matrices are estimated after residualizing the data on age, sex, and diagnosis status. Scanner A is a Siemens Symphony 1.5T scanner with 23 subjects and the other scanners are General Electric Signa Excite 1.5T scanners with 20 subjects each.

**Table 1:** ADNI demographics by scanner for the three scanners with the largest number of acquired subjects. Manufacturer of each scanner is displayed in parentheses. ANOVA p-values for testing differences in the mean of continuous variables and Chi-squared test p-values for testing the differences in categorical variables are reported in the rightmost column.

|  | A (Siemens) | B (General Electric) | C (General Electric) | *p* |
| --- | --- | --- | --- | --- |
| Number of Subjects | 23 | 20 | 20 |  |
| Age (mean (SD)) | 74.48 (5.13) | 76.90 (8.18) | 78.78 (6.19) | 0.11 |
| Diagnosis (%) |  |  |  | 0.57 |
| AD | 7 (30.4) | 6 (30.0) | 5 (25.0) |  |
| CN | 6 (26.1) | 5 (25.0) | 2 (10.0) |  |
| LMCI | 10 (43.5) | 9 (45.0) | 13 (65.0) |  |
| Male (%) | 10 (43.5) | 13 (65.0) | 16 (80.0) | 0.05 |

We also provide quantitative comparisons for pairwise distances across scanners before and after harmonization in Table 2. A tuning parameter of the CovBat model is the desired proportion of variance explained in the dimension reduction space, which we select at 95% (23 PCs). To ensure that our results do not depend strongly on the choice of tuning parameter, we also report the minimum and maximum of the pairwise Frobenius norms after applying CovBat with percent variation explained ranging from 56% (2 PCs) to 100% (62 PCs). We report the results of this sensitivity analysis in parentheses. We find that ComBat adjustment can modestly harmonize the covariance matrices but CovBat adjustment shows large reductions in the between-scanner distances across a range of tuning parameter choices.

**Table 2:** Pairwise distances between scanner-specific covariance matrices. Differences in covariance structure between scanner are reported as the Frobenius distance between covariance matrices calculated across observations acquired on each scanner. Results from adjusting the number of PC scores ranging from those explaining 56% to 100% of variation are shown in parentheses as the minimum and maximum pairwise Frobenius norms across the range.

|  | Unharmonized | ComBat | CovBat |
| --- | --- | --- | --- |
| A,B | 6.28 | 5.26 | 3.59 (3.58 - 3.69) |
| A,C | 5.94 | 4.59 | 3.25 (3.09 - 3.26) |
| B,C | 4.32 | 3.76 | 3.42 (3.42 - 3.81) |

### 3.2 CovBat impairs detection of scanner

To evaluate the potential impact of scanner effects in covariance using MVPA, we conduct a Monte-Carlo split-sample experiment for prediction of scanner manufacturer labels using all 213 ADNI subjects before and after harmonization with existing methods. We train using data harmonized with the state-of-the-art ComBat method and our proposed method, CovBat. Fig. 3a shows that Siemens scanners are easily identifiable based on unharmonized cortical thickness measurements (median area-under-the-curve (AUC) 0.89, IQR 0.87-0.90), which is consistent with recent findings (Glocker et al., 2019). We also note that scanner manufacturer is still detected after ComBat is applied (0.66, 0.64-0.68). After CovBat, the machine learning method’s performance for differentiating between scanners is close to chance (0.46, 0.44-0.48). DeLong’s test results shown in **Supplementary Figure 1** suggest that these AUC values are significantly different between ComBat and CovBat. Using MANOVA, Table 3 shows that the association with scanner manufacturer is statistically significant in unharmonized and ComBat-adjusted data but is eliminated in CovBat-adjusted data.

**Figure 3:**
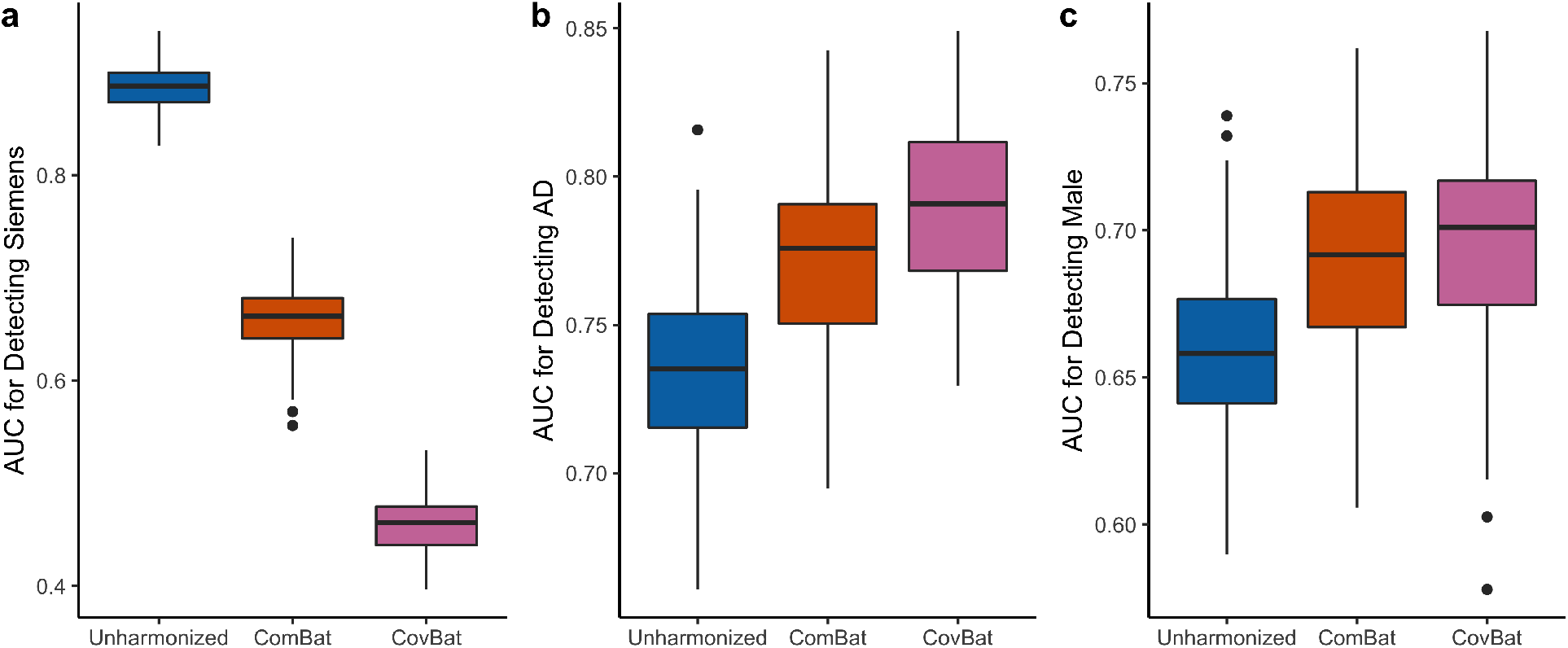
Multivariate pattern analysis experiments for detection of scanner manufacturer, sex, and Alzheimer’s disease status using cortical thickness data. The data is randomly split into 50% training and 50% validation then used to train a random forests algorithm to predict a specified trait. AUC values from 100 repetitions of this analysis are reported for unharmonized, ComBat-adjusted, and CovBat-adjusted data. **a**, Boxplot showing results for detecting if subjects were acquired on a Siemens scanner. Results for detection of Alzheimer’s disease status are shown in **b** and results for detection of sex are shown in **c**.

**Table 3:** Multivariate analysis of variance *p*-values for scanner manufacturer, sex, and Alzheimer’s disease status using cortical thickness data. The analysis is performed using Pillai’s trace as the test statistic.

|  | Unharmonized | ComBat | CovBat |
| --- | --- | --- | --- |
| Manufacturer | < 0.001 | < 0.001 | 1 |
| Sex | < 0.001 | < 0.001 | < 0.001 |
| Diagnosis | < 0.001 | < 0.001 | < 0.001 |

### 3.3 CovBat retains biological associations

It is well-known that cortical thickness differs substantially by sex and Alzheimer’s disease status (Lerch et al., 2005; Sowell et al., 2007). To assess whether CovBat maintains biological associations of interest, we perform two MVPA experiments using the full ADNI data to classify healthy versus Alzheimer’s disease (AD) and to differentiate patients by sex. Fig. 3b,c show that both of these biological associations are retained after either harmonization method. For AD classification, the median AUC increases from 0.74 (IQR 0.72 - 0.75) in unharmonized data to 0.78 (0.75 - 0.79) in ComBat-adjusted data to 0.79 (0.77-0.81) in CovBat-adjusted. Similarly, the median AUC for detection of sex increased from 0.66 (0.64 - 0.68) to 0.69 (0.67 - 0.71) to 0.70 (0.67 - 0.72). For detection of both AD and male sex, DeLong’s test results plotted in **Supplementary Figure 1** support that the AUCs are not significantly different between ComBat and CovBat. These findings suggest that CovBat not only provides thorough removal of scanner effects, but also maintains clinical associations. **Appendix A.1** shows that similar results hold for prediction of age. **Appendix A.2** shows that these results for both detection of site and biological associations largely hold even when CovBat is trained on a subset of the data. MANOVA results in Table 3 show that the significant associations with diagnosis and sex are retained after either harmonization method.

### 3.4 Simulation studies

We first perform simulations to assess whether either harmonization method can recover the underlying covariance structure in our simple simulation setting and harmonize covariance matrices generally. In the Simple Covariance Effects simulation, Fig. 4 shows that CovBat outperforms ComBat in recovery of the true covariance structure (denoted *S* in **Section 2.5.1**) across all parameters considered. Remaining deviation from the true covariance can be explained by error in covariance estimation; even with 10000 samples per site and 62 features the distance of the pooled covariance estimate from the true covariance is still 3.14. In Table 4, we observe in the same simulation setting with a within-site sample size of 250 and 62 simulated features that CovBat performs best in harmonizing within-site covariance matrices. That result is replicated with a more complex covariance site effect in the Predictor Affects Mean simulation as shown in Table 5.

**Figure 4:**
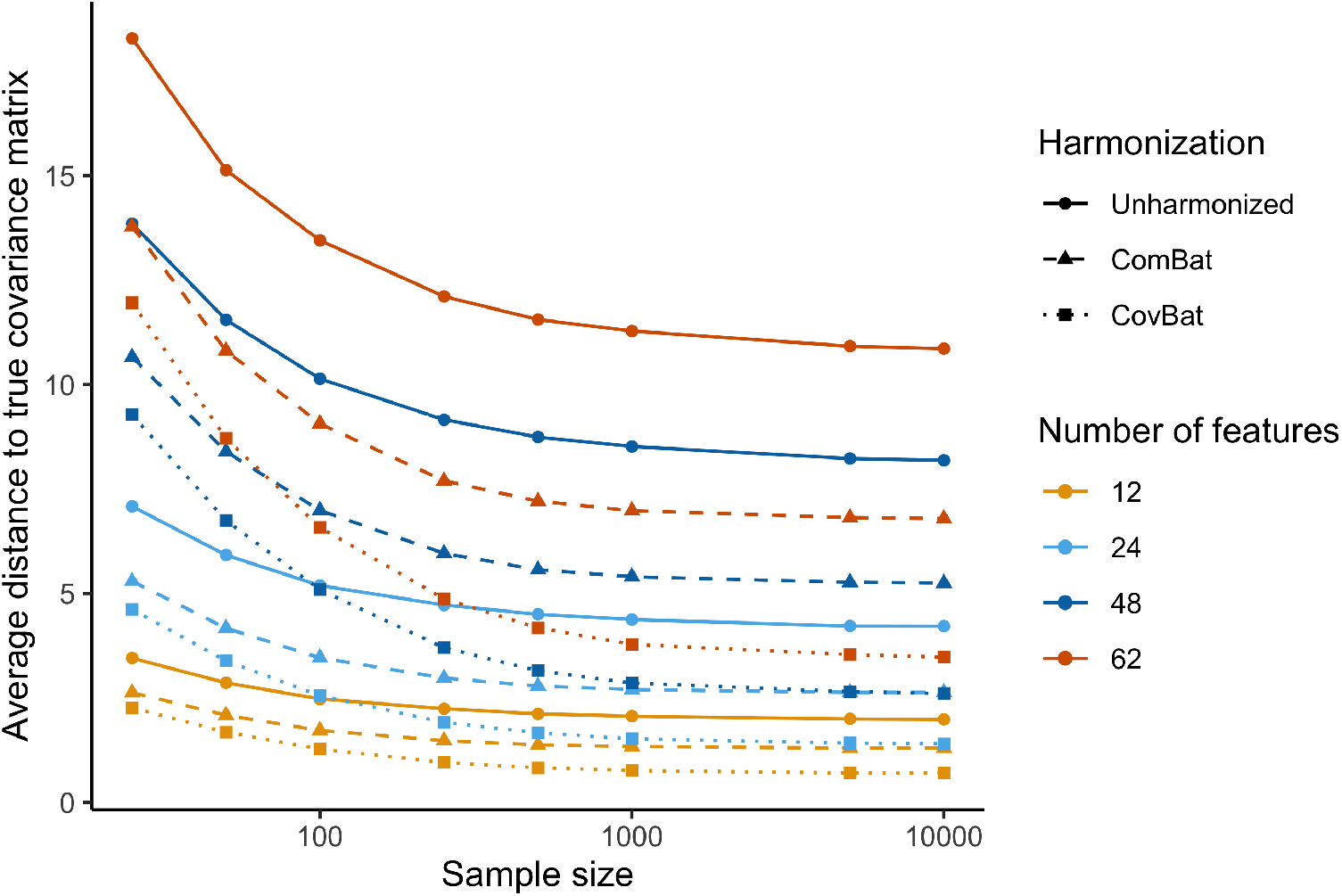
Average across sites of the Frobenius distance between sample site-specific covariance matrices and the true covariance matrix for the Simple Covariance Effects simulation. The displayed values are averaged across the mean Frobenius distance for each site, which are taken across 1000 simulations each. Results are plotted for a sample size per site of 25, 50, 100, 250, 500, 1000, 5000, and 10000.

**Table 4:**
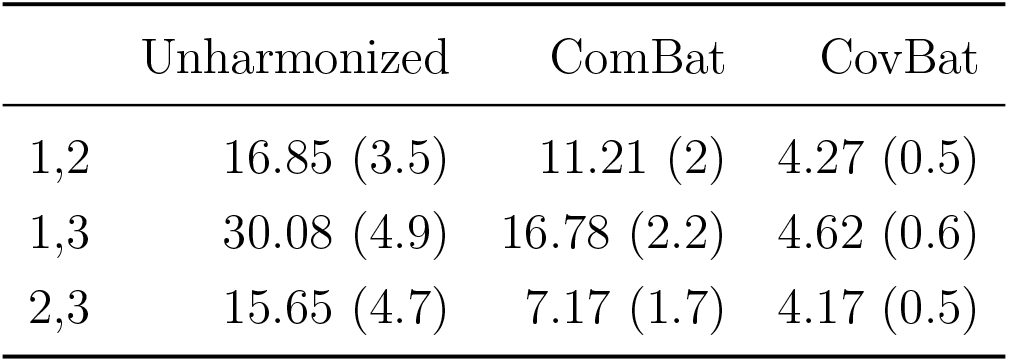
Mean and standard deviation of pairwise Frobenius norms between within-site covariance estimates for the Simple Covariance Effects simulation. Standard deviations are reported in parentheses.

|  | Unharmonized | ComBat | CovBat |
| --- | --- | --- | --- |
| 1,2 | 16.85 (3.5) | 11.21 (2) | 4.27 (0.5) |
| 1,3 | 30.08 (4.9) | 16.78 (2.2) | 4.62 (0.6) |
| 2,3 | 15.65 (4.7) | 7.17 (1.7) | 4.17 (0.5) |

**Table 5:** Mean and standard deviation of pairwise Frobenius norms between within-site covariance estimates for the Predictor Affects Mean simulation. Standard deviations are reported in parentheses.

|  | Unharmonized | ComBat | CovBat |
| --- | --- | --- | --- |
| 1,2 | 15.77 (2.1) | 15.39 (1.4) | 12.81 (0.8) |
| 1,3 | 13.22 (1.4) | 13.68 (1.3) | 11.69 (0.7) |
| 2,3 | 12.79 (1.8) | 10.98 (0.7) | 10.98 (0.5) |

We then evaluate CovBat through two MVPA experiments across all simulation settings considered. For our main analyses, we simulate 62 cortical thicknesses for 250 subjects per site. We begin by examining a ComBat simulation, where we generate data from the original ComBat model. That is, we impose mean and variance site effects while also simulating a predictor that has an effect on the mean. Fig. 5a,b show that CovBat performs almost identically to ComBat in this scenario, showing that our method performs competitively in the absence of covariance effects. In the Predictor Affects Mean simulation, Fig. 5c,d shows that CovBat also substantially reduces the chance of detecting site and performs similarly to ComBat for detection of the predictor. These simulations demonstrate that CovBat performs at least as well or better than ComBat when the predictor affects only the mean of the observations.

**Figure 5:**
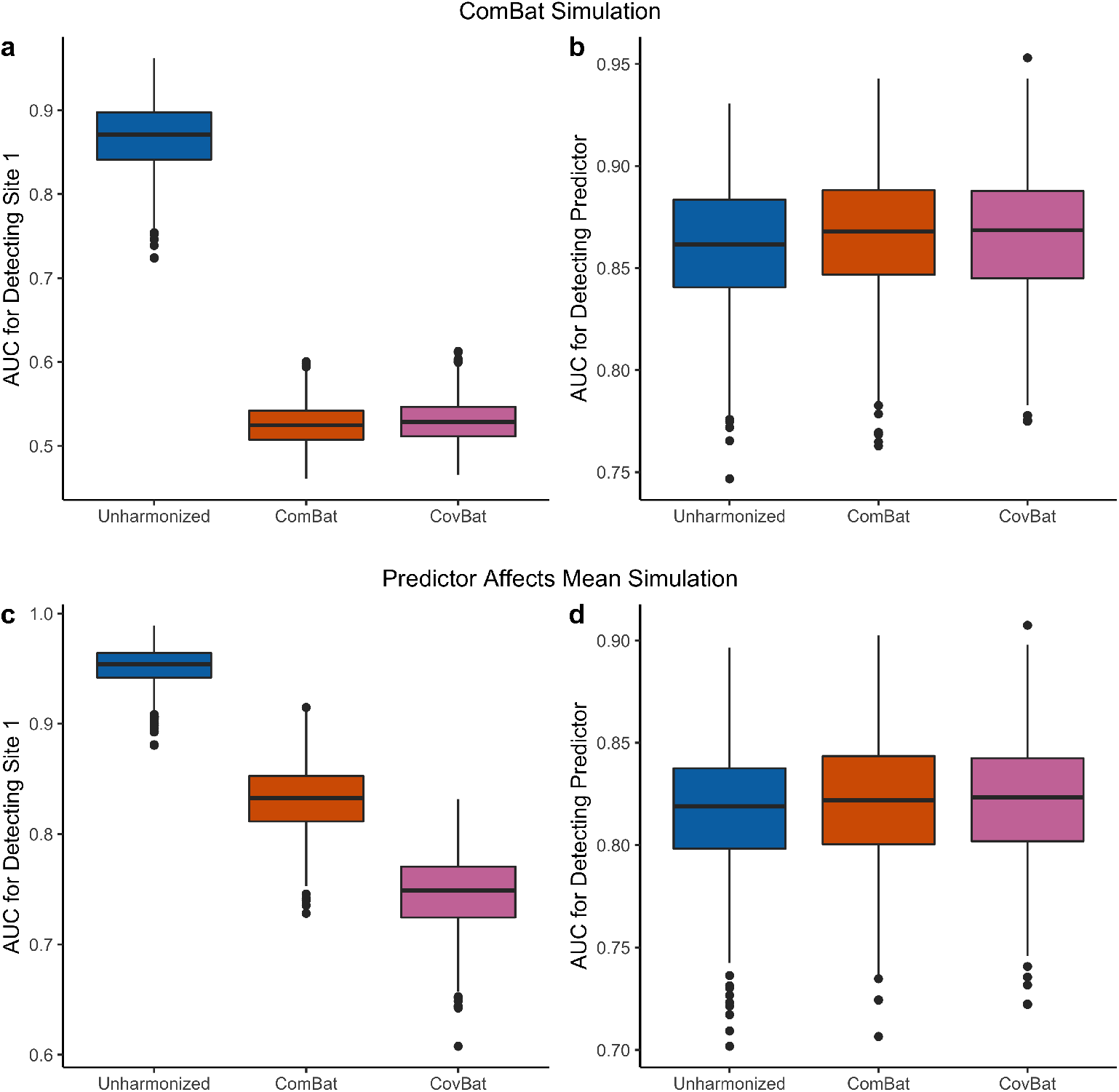
Results from MVPA simulations for detection of site and for detection of a binary outcome in the absence of predictor effects on covariance. The simulated data consists of 62 cortical thicknesses for 250 subjects per site across three sites. For each of 1000 simulations, the data is randomly split into 50% training and 50% validation. A random forests algorithm is trained using the training set to predict either Site 1 or the presence of the binary outcome. **a**, Boxplot showing Site 1 detection in the ComBat simulation. **b**, Boxplot showing predictor detection in the ComBat simulation. **c**, Boxplot showing Site 1 detection in the Predictor Affects Mean simulation. **d**, Boxplot showing predictor detection in the Predictor Affects Mean simulation.

We then incorporate a predictor effect on covariance in our Predictor Affects Covariance simulation and show that CovBat reduces detection of site (Fig. 6a) and maintains the association with the predictor (Fig. 6b). In order to further emphasize the importance of covariance effects, we investigate a Covariance Only simulation where both site and predictor effects exist only in the covariance of observations, but not the mean or variance. In unharmonized data, we observe high mean AUC values for detection of Site 1 and detection of the predictor, both of which are essentially unaffected after implementing ComBat. After CovBat though, we see substantial improvements on both metrics (Fig. 6c,d). While CovBat performs well in both simulations, we find that CovBat does not entirely remove the severe covariance site effect. Regardless, we observe that CovBat offers notable improvements over ComBat owing to its harmonization of covariance.

**Figure 6:**
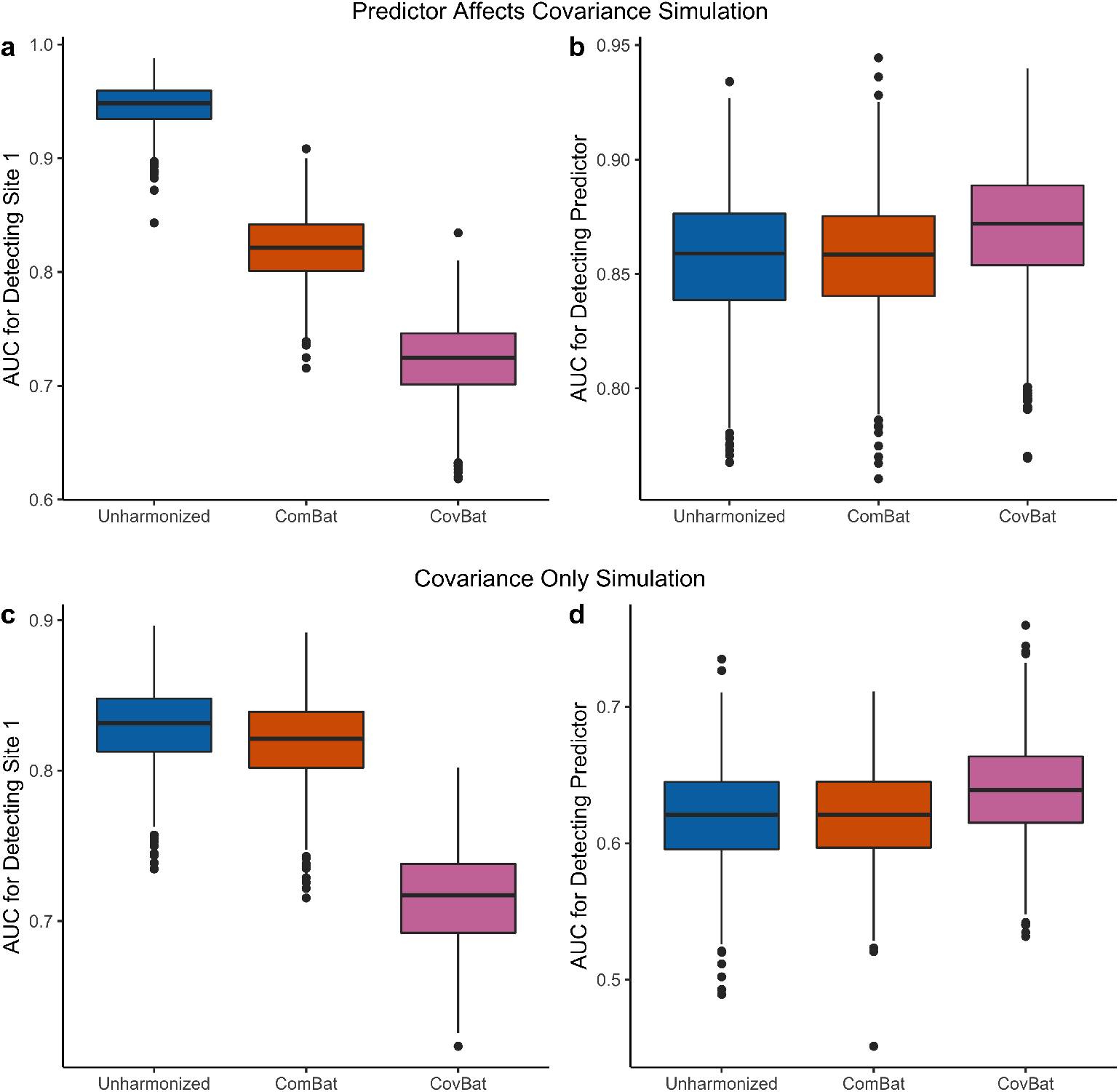
Results from MVPA simulations for detection of site and for detection of a binary clinical outcome where the predictor also confounds covariance. The simulated data consists of 62 cortical thicknesses for 250 subjects per site across three sites. For each of 1000 simulations, the data is randomly split into 50% training and 50% validation. A random forests algorithm is trained using the training set to predict either Site 1 or the presence of the binary predictor. **a**, Boxplot showing Site 1 detection in the Predictor Affects Covariance simulation. **b**, Boxplot showing predictor detection in the Predictor Affects Covariance simulation. **c**, Boxplot showing Site 1 detection in the Covariance Only simulation. **d**, Boxplot showing predictor detection in the Covariance Only simulation.

We then conduct additional analyses to assess the robustness of CovBat to reductions in sample size per site and number of features. In **Supplementary Table 1**, we show that the random forests MVPA results largely hold for simulations without a predictor effect on covariance; however, in the absence of a covariance site effect CovBat performs slightly worse overall and in situations with small sample size per site (*n_i_* = 25) and larger number of features (*p* ≥ 48) we observe that CovBat can inflate detection of site. **Supplementary Table 2** shows that similar results hold in simulations with a predictor effect on covariance with CovBat showing good performance overall but poor performance with small sample sizes and large number of features. To assess if these negative findings may be tied to the random forests implementation of MVPA, we additionally perform MANOVA for site and diagnosis status across Predictor Affects Mean and Predictor Affects Covariance simulations and show in **Supplementary Table 3** and **Supplementary Table 4** that associations with site are reduced after CovBat across all scenarios while associations with diagnosis are preserved. We repeat these MANOVA analyses in the Simple Covariance Effects simulation and show in **Supplementary Table 5** that CovBat performs better than ComBat at controlling the rejection rate across nearly all parameters considered. However, CovBat only successfully controls the rejection rate at 5% in simulations with less features, small sample size, and large sample size. The controlled rejection rate in small samples however could be explained by the low power of MANOVA in small sample size relative to the number of features (Stevens, 1980).

## 4 Discussion

The growing number of multi-site studies across diverse fields has spurred the development of harmonization methods that are general, but also account for field-specific challenges. In neuroimaging research, the rise of MVPA has established an unmet need for harmonization of covariance. We demonstrate that strong scanner effects in covariance exist, influence downstream MVPA experiments, and remain after performing the state-of-the-art harmonization. We then propose a novel method and demonstrate that it is effective in removing scanner differences in covariance and retaining the detection of biological associations via MVPA. Simulation studies show similar MVPA results and demonstrate that CovBat performs well across a variety of settings and sample sizes.

In ADNI data, we show that substantial differences exist in the covariance structures of cortical thickness observations and can be mitigated through our proposed method. We furthermore show that MVPA can detect these scanner effects, whether through machine learning or conventional multivariate analyses. These results mirror recent studies that predict scanner from neuroimaging features with high accuracy (Glocker et al., 2019) and a recent study demonstrating that ComBat is insufficient to prevent detection of Siemens-manufactured scanners in a large multi-site dataset(Nielson et al., 2018). We then demonstrate that CovBat can almost entirely prevent scanner detection in the ADNI dataset. To ensure generalizability of these results, implementation of CovBat in other multi-site studies of varying experimental designs should be pursued in the future.

In simulation, CovBat shows generally strong performance in removing scanner effects in medium and large sample sizes across varying number of features and complexity of the scanner effect. We demonstrate that CovBat almost fully removes scanner effects when they exist in the principal component directions; however, MVPA results show that scanner effects may remain in more complex scenarios. Caution should be taken in attempting to address covariance scanner effects in smaller samples with many features. We show potential increases in scanner detection in these situations which are potentially the result of poor covariance estimation in high-dimensional settings. While we do not observe these limitations through MANOVA, we also acknowledge that MANOVA may be underpowered in our scenarios with low sample size and high dimensionality as shown in previous studies (Stevens, 1980).

Our proposed method harmonizes covariance across scanners by removing mean and variance shifts in the the principal components space, which we show to be effective in addressing the covariance effects we observe. This idea resembles spectral models which also relate covariates to the eigendecomposition of covariance matrices (Boik, 2002). Our method assumes that the ideal covariance structure exists in the eigenspace of the full data covariance matrix. As we show through our simulations, this model may be insufficient to remove scanner effects which do not resemble the covariance structure of the data. Potential extensions could incorporate methods that model scanner effects as separate low-rank structures (Hoff & Niu, 2012) or identify projections most related to scanner (Zhao et al., 2019). However, implementation of these methods in a harmonization framework may not be as straightforward as our proposed method.

A limitation of our methodology is that CovBat is a covariate-assisted harmonization method similar to ComBat and requires specification of covariates to protect in the data. Associations with covariates not included in the harmonization step can certainly be removed alongside scanner effects and recent papers have even identified situations where spurious associations can be introduced (Nygaard et al., 2016; Zindler et al., 2020). While we do not observe CovBat introducing false positives in our investigation, care must be taken in implementing CovBat protecting for the outcome of interest especially in unbalanced study designs. We reiterate previous advice that analyses should be performed with and without harmonization and that associations detected only after harmonization should be interpreted with caution (Zindler et al., 2020).

Our study demonstrates that scanner effects can exist in the covariance of structural neuroimaging data and can be mitigated via our proposed methodology. Future studies should determine how scanner properties can influence the covariance structure of the data and if other multi-site multivariate neuroimaging studies contain similar effects. Further methodological work could utilize other covariance modeling strategies in order to address more complex scanner effects. Since our method operates on general multivariate data, our findings extend directly to functional, metabolic, and other imaging modalities. Further studies should also determine the extent to which multivariate statistical and machine learning studies of genomic data are susceptible to the biases documented.

## 5 Software

All of the postprocessing analysis was performed in the R statistical software (V3.6.1). CovBat is available for both R and Python (https://github.com/andy1764/CovBat_Harmonization). Reference implementations for ComBat are available in R and Matlab (https://github.com/Jfortin1/ComBatHarmonization) and in Python (https://github.com/ncullen93/neuroCombat).

## 6 Acknowledgements

This work was supported by the National Institute of Neurological Disorders and Stroke (grant numbers R01 NS085211 and R01 NS060910), the National Multiple Sclerosis Society (RG-1707-28586) and a seed grant from the University of Pennsylvania Center for Biomedical Image Computing and Analytics (CBICA). The content is solely the responsibility of the authors and does not necessarily represent the official views of the funding agencies.

The majority of the data used in this paper are derived from the ADNI study. Data collection and sharing for this project was funded by the Alzheimer’s Disease Neuroimaging Initiative (ADNI) (National Institutes of Health Grant U01 AG024904) and DOD ADNI (Department of Defense award number W81XWH-12-2-0012). ADNI is funded by the National Institute on Aging, the National Institute of Biomedical Imaging and Bioengineering, and through generous contributions from the following: AbbVie, Alzheimer’s Association; Alzheimer’s Drug Discovery Foundation; Araclon Biotech; BioClinica, Inc.; Biogen; Bristol-Myers Squibb Company; CereSpir, Inc.; Cogstate; Eisai Inc.; Elan Pharmaceuticals, Inc.; Eli Lilly and Company; EuroImmun; F. Hoffmann-La Roche Ltd and its affiliated company Genentech, Inc.; Fujirebio; GE Healthcare; IXICO Ltd.;Janssen Alzheimer Immunotherapy Research & Development, LLC.; Johnson & Johnson Pharmaceutical Research & Development LLC.; Lumosity; Lundbeck; Merck & Co., Inc.;Meso Scale Diagnostics, LLC.; NeuroRx Research; Neurotrack Technologies; Novartis Pharmaceuticals Corporation; Pfizer Inc.; Piramal Imaging; Servier; Takeda Pharmaceutical Company; and Transition Therapeutics. The Canadian Institutes of Health Research is providing funds to support ADNI clinical sites in Canada. Private sector contributions are facilitated by the Foundation for the National Institutes of Health (www.fnih.org). The grantee organization is the Northern California Institute for Research and Education, and the study is coordinated by the Alzheimer’s Therapeutic Research Institute at the University of Southern California. ADNI data are disseminated by the Laboratory for Neuro Imaging at the University of Southern California.

## Competing interests

The authors declare no competing interests.

## Author contributions

R.T.S. and H.S. contributed equally to this work. **Andrew A. Chen:** Formal analysis, Conceptualization, Methodology, Software, Investigation, Writing - original Draft **Joanne C. Beer:** Data curation, Writing - review & editing **Nicholas J. Tustison:** Data curation, Writing - review & editing **Philip A. Cook:** Data curation, Writing - review & editing **Russell T. Shinohara:** Supervision, Conceptualization, Methodology, Writing - review & editing **Haochang Shou:** Supervision, Conceptualization, Methodology, Writing - review & editing

## Appendix A

## A.1 CovBat preserves prediction of age

ComBat has previously been shown to preserve age prediction via linear regression and support vector regression (Fortin et al., 2018). To evaluate whether this result holds in ADNI data for ComBat or CovBat, we propose an additional MVPA experiment. We i) randomly split the subjects into 50% training set and 50% validation set, ii) train a random forests algorithm to predict age, and iii) assess predictive performance on the validation set via root-mean-square-error (RMSE). We perform steps (i)-(iii) 100 times each for unharmonized, ComBat-adjusted, and CovBat-adjusted data. Fig. A.1. shows that the mean RMSE for age prediction decreases from 6.05 (±0.22) in unharmonized to 5.81 (±0.23) in ComBat-adjusted data to 5.75 (±0.21) in CovBat-adjusted data. These findings are consistent with previous work (Fortin et al., 2018) and show that CovBat provides similar recovery of the association between cortical thickness and age.

## A.2 Parameter estimation using training subset

Both ComBat and CovBat estimate and residualize out the covariate effects using the full data; however, there are cases were only a subset of the data is available when performing harmonization. For instance, if a group of subjects has already been acquired, prediction on subjects subsequently acquired on the same scanners could only leverage data from the original sample. In this scenario, the new sample can be harmonized using ComBat or CovBat by estimating the covariate effect using the original sample, then proceeding with subsequent steps as usual.

**Figure A.1.:**
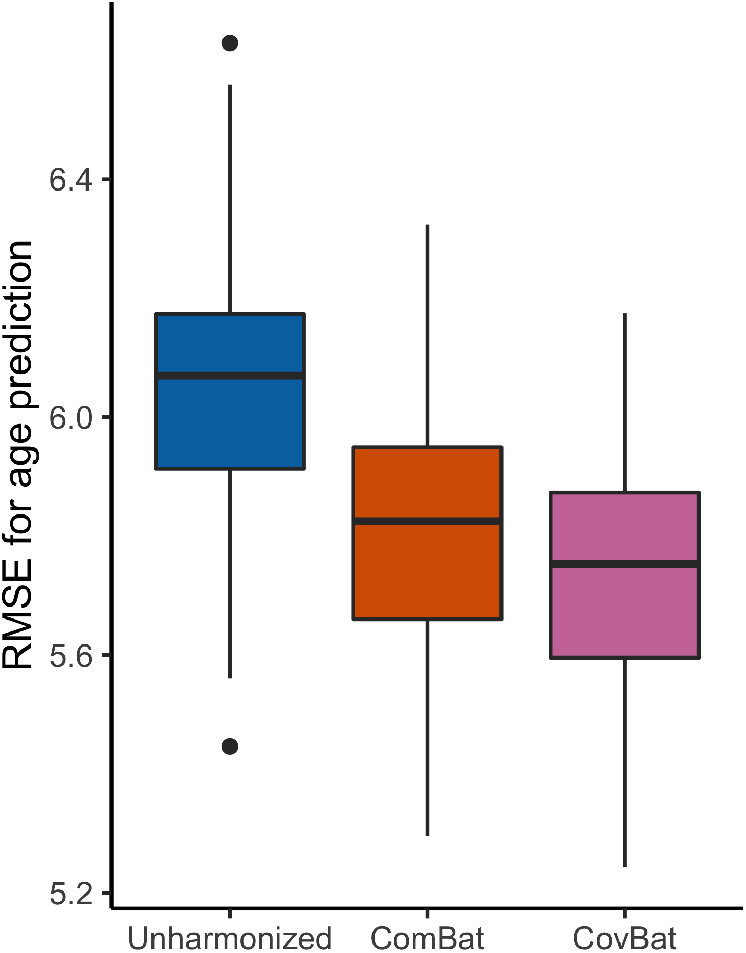
Multivariate pattern analysis experiments for detection of age using cortical thickness data. The data is randomly split into 50% training and 50% validation then used to train a random forests algorithm to predict age. RMSE values from 100 repetitions of this analysis are reported for unharmonized, ComBat-adjusted, and CovBat-adjusted data.

We evaluate this modification by repeating our main MVPA analyses using ADNI data with different subsampling of the patients. Specifically, we replace step (i) in our MVPA experiments by instead splitting the sample into 270 training subjects and 235 testing subjects such that both the train and test sets contain at least one subject acquired on each scanner. We then apply ComBat and CovBat by estimating the covariate effects using only the training subjects. Fig. A.2. shows the results for all MVPA experiments. Detection of site (AUC 0.89 ± 0.02 in raw data) still worsens after ComBat (0.66 ± 0.03) and is almost at chance after CovBat (0.54 ± 0.03). For detection of AD, improvements over unharmonized (AUC 0.74 ± 0.03) are still demonstrated after ComBat adjustment (0.77 ± 0.03) and CovBat adjustment (0.78 ± 0.02). For detection of male, lesser improvement is observed from unharmonized (AUC 0.67 ± 0.03) to ComBat (0.68 ± 0.03) to CovBat (0.68 ± 0.03). Mean RMSE for age prediction decreases from 5.99 (±0.22) in unharmonized to 5.86 (±0.23) in ComBat-adjusted data to 5.82 (±0.22) in CovBat-adjusted data. Overall, the results appear quite similar to harmonization using the full dataset, showing that CovBat performs well even when only a limited training subset is available.

**Figure A.2.:**
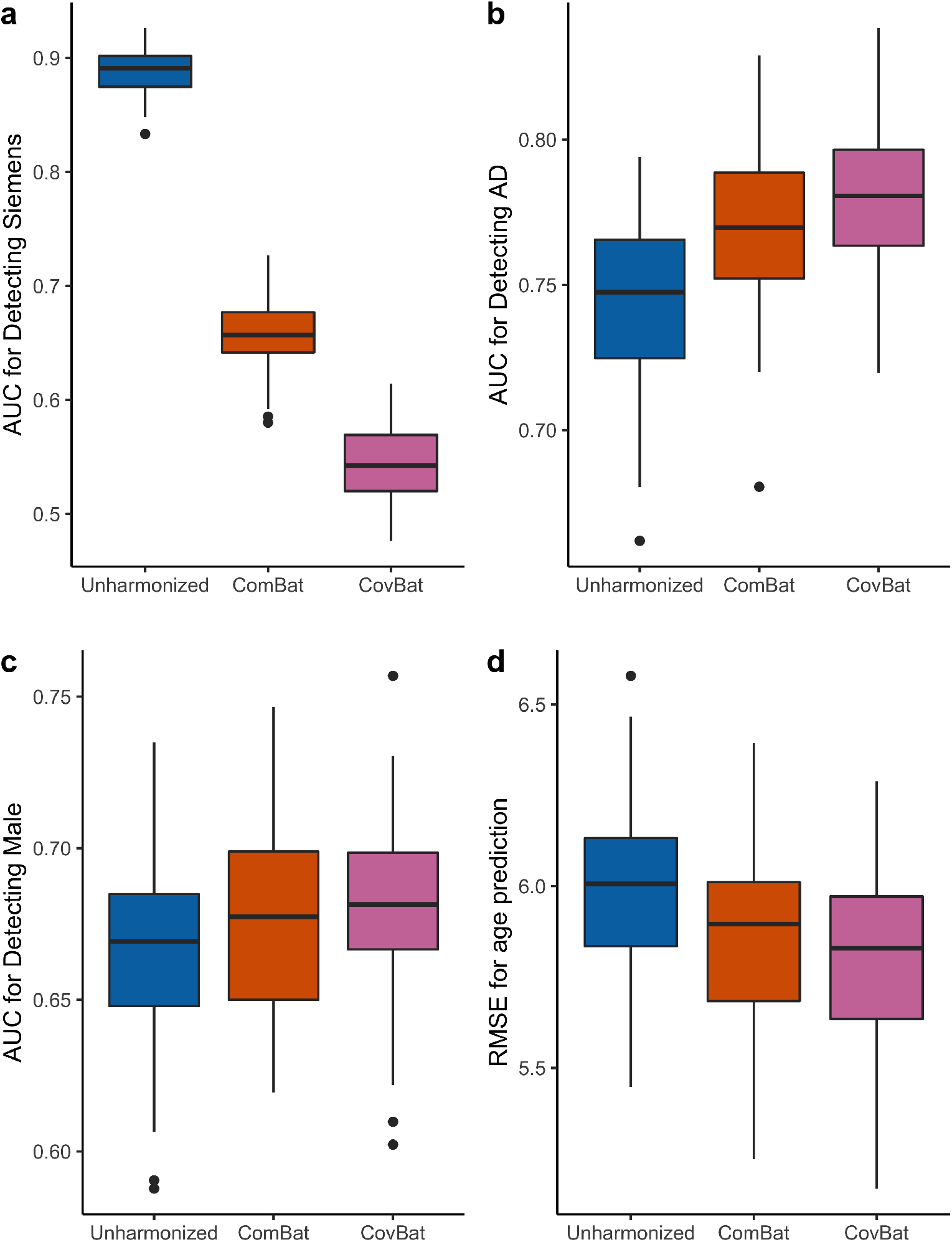
MVPA experiment results for harmonization using only training data. The data is randomly split into 270 training subjects and 235 testing subjects such that every site is represented in each group. The training set is then used to train a random forests algorithm to predict scanner or patient characteristics. **a** shows the AUC values for detection of site. AUC values for detection of AD are displayed in **b** and detection of male in **c**. RMSE values for prediction of age are displayed in **d**.

## Notes

### Competing Interest Statement

The authors have declared no competing interest.

### Summary of Updates

New work undertaken -To evaluate a more classical analysis technique, we conducted multivariate analysis of variance (MANOVA) as an alternative approach of multivariate pattern analysis (MVPA) and show that mitigation of scanner effects and preservation of biological variability are achieved. -We conduct signed-rank tests to evaluate differences in AUC in the MVPA experiment. -We designed a new simulation to evaluate how our proposed method performs in a scenario with simple covariance site effects for which the CovBat model is well specified. -In this new Simple Covariance Effects simulation, we compare sample site-specific covariance matrices to true covariance without site effects. These analyses show the degree of error introduced by covariance estimation and site confounding before and after harmonization. -We have conducted additional simulations to evaluate robustness of the MVPA results to sample size and number of features. Major changes to the manuscript -The introduction includes additional discussions and details on why harmonization using neither generative adversarial networks (GANs) nor a distance-based approach are both not directly comparable to our proposed method. -Metrics based on correlation have been modified to instead focus on covariance. -Section 2.1 has been clarified to better explain the two samples that are used throughout. -Section 2.1 includes a brief statement to explain why the harmonization method proposed in Zhou et al. (2018) cannot apply to the ADNI dataset used throughout the paper. -Section 2.3 now includes more detail as to the rationale and implementation of CovBat. -Section 2.5 has been reworked to describe two different simulation designs and multiple scenarios among those designs. -Section 3.2 and 3.3 include MANOVA results for scanner manufacturer, sex, and diagnosis. -Section 3.4 includes a figure to show that harmonization better recovers the true covariance structure in the Simple Covariance Effects simulation. -The Discussion section has been expanded to comment on all major results on the paper and acknowledge limitations of the methodology and study design. Minor edits and fixes -The description of the sensitivity analysis in Section 3.1 now matches the caption of Table 2. -Captions have been reduced in length and we no longer report mean and standard deviation, instead opting to report median and interquartile range to be more consistent with the markers shown on the plots.

