## Supplementary materials for "Removal of Scanner Effects in Covariance Improves Multivariate Pattern Analysis in Neuroimaging Data"

[e]Data used in preparation of this article were obtained from the Alzheimer's Disease Neuroimaging Initiative (ADNI) database (adni.loni.usc.edu). As such, the investigators within the ADNI contributed to the design and implementation of ADNI and/or provided data but did not participate in analysis or writing of this report. A complete listing of ADNI investigators can be found at: http://adni.loni.usc.edu/wp-content/uploads/how_to_apply/ADNI_Acknowledgement_List.pdf

[1]Equal contribution

***Correspondence: Andrew A. Chen**

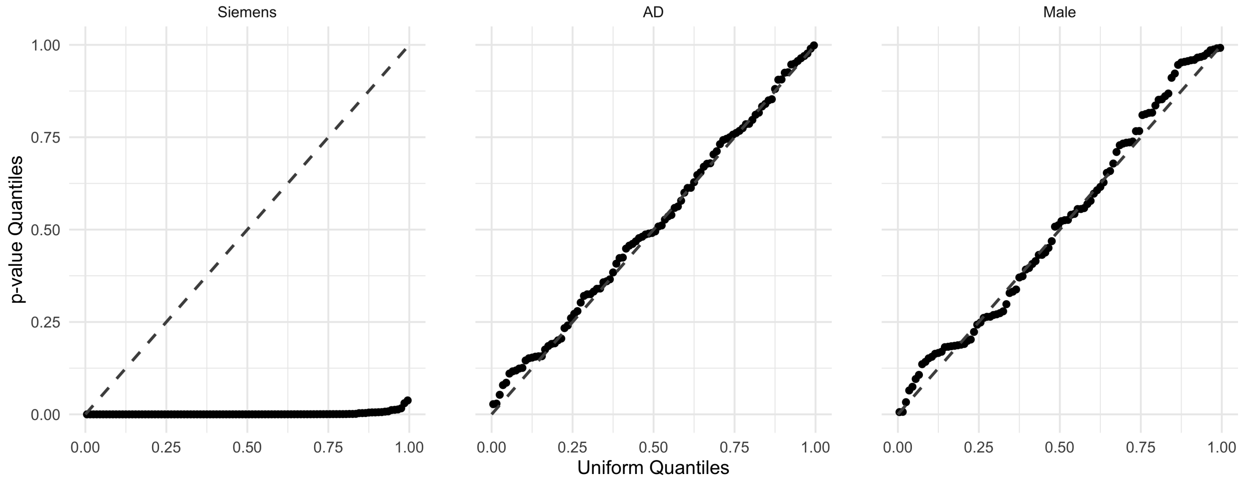


Supplementary Figure 1: **DeLong's test $p$-value Q-Q plots for comparing AUC between ComBat and CovBat.** For each of the 100 train-test splits within each MVPA experiment, DeLong's test is performed with the two-sided null hypothesis that the AUC is different in ComBat-adjusted and CovBat-adjusted data. The $p$-values for each experiment are compared to a uniform distribution from 0 to 1.

| | | ComBat Simulation | | | Predictor Affects Mean | | |
|---|---|---|---|---|---|---|---|
| $n_i$ | $p$ | Unharmonized | ComBat | CovBat | Unharmonized | ComBat | CovBat |
| 25 | 12 | 0.61 (0.54-0.67) | 0.58 (0.53-0.63) | 0.62 (0.56-0.67) | 0.69 (0.63-0.76) | 0.58 (0.53-0.63) | 0.57 (0.52-0.62) |
| | 24 | 0.64 (0.57-0.71) | 0.58 (0.53-0.63) | 0.64 (0.58-0.69) | 0.74 (0.68-0.80) | 0.58 (0.53-0.65) | 0.56 (0.52-0.61) |
| | 48 | 0.67 (0.60-0.74) | 0.58 (0.53-0.63) | 0.66 (0.61-0.71) | 0.72 (0.65-0.78) | 0.56 (0.52-0.61) | 0.62 (0.57-0.68) |
| | 62 | 0.68 (0.61-0.75) | 0.58 (0.53-0.63) | 0.66 (0.61-0.72) | 0.72 (0.65-0.79) | 0.56 (0.52-0.61) | 0.64 (0.58-0.70) |
| 50 | 12 | 0.64 (0.58-0.70) | 0.55 (0.52-0.59) | 0.58 (0.53-0.62) | 0.79 (0.74-0.83) | 0.68 (0.62-0.73) | 0.59 (0.55-0.64) |
| | 24 | 0.69 (0.64-0.75) | 0.55 (0.52-0.59) | 0.59 (0.54-0.63) | 0.84 (0.80-0.88) | 0.72 (0.67-0.78) | 0.59 (0.55-0.64) |
| | 48 | 0.72 (0.67-0.78) | 0.55 (0.52-0.59) | 0.61 (0.57-0.65) | 0.82 (0.77-0.86) | 0.61 (0.56-0.67) | 0.54 (0.51-0.57) |
| | 62 | 0.74 (0.69-0.79) | 0.55 (0.52-0.59) | 0.61 (0.57-0.65) | 0.81 (0.77-0.86) | 0.58 (0.54-0.63) | 0.55 (0.51-0.58) |
| 100 | 12 | 0.67 (0.63-0.72) | 0.54 (0.51-0.56) | 0.55 (0.52-0.58) | 0.86 (0.83-0.89) | 0.78 (0.75-0.82) | 0.71 (0.68-0.75) |
| | 24 | 0.74 (0.70-0.78) | 0.54 (0.51-0.56) | 0.56 (0.53-0.59) | 0.92 (0.90-0.94) | 0.85 (0.82-0.88) | 0.74 (0.71-0.78) |
| | 48 | 0.79 (0.75-0.83) | 0.54 (0.51-0.57) | 0.57 (0.54-0.60) | 0.90 (0.87-0.93) | 0.75 (0.71-0.78) | 0.62 (0.58-0.66) |
| | 62 | 0.82 (0.78-0.85) | 0.54 (0.51-0.57) | 0.57 (0.54-0.60) | 0.90 (0.87-0.93) | 0.71 (0.67-0.76) | 0.60 (0.57-0.64) |
| 250 | 12 | 0.72 (0.67-0.75) | 0.52 (0.51-0.54) | 0.53 (0.51-0.55) | 0.92 (0.91-0.94) | 0.88 (0.87-0.90) | 0.84 (0.83-0.86) |
| | 24 | 0.80 (0.77-0.84) | 0.52 (0.51-0.54) | 0.53 (0.51-0.55) | 0.97 (0.96-0.98) | 0.95 (0.94-0.96) | 0.90 (0.88-0.92) |
| | 48 | 0.87 (0.84-0.89) | 0.53 (0.51-0.55) | 0.54 (0.52-0.56) | 0.96 (0.95-0.97) | 0.88 (0.87-0.90) | 0.79 (0.77-0.81) |
| | 62 | 0.89 (0.86-0.91) | 0.53 (0.51-0.55) | 0.54 (0.52-0.56) | 0.97 (0.96-0.97) | 0.87 (0.85-0.88) | 0.79 (0.76-0.81) |

Supplementary Table 1: **AUC results from MVPA simulations for detection of site across multiple sample sizes (n) and number of features (p) in the absence of predictor effects on covariance.** For each of 1000 simulations, the data is randomly split into 50% training and 50% validation then a random forests algorithm is trained using the training set to predict either Site 1 or the presence of the binary predictor. The median AUC across these simulations are reported with lower and upper quartiles displayed in parentheses. Scenarios where CovBat outperforms ComBat are colored in blue.

| | | Predictor Affects Covariance | | | Covariance Only | | |
|---|---|---|---|---|---|---|---|
| $n_i$ | $p$ | Unharmonized | ComBat | CovBat | Unharmonized | ComBat | CovBat |
| 25 | 12 | 0.70 (0.63-0.77) | 0.58 (0.53-0.63) | 0.57 (0.53-0.62) | 0.61 (0.56-0.68) | 0.58 (0.53-0.63) | 0.57 (0.52-0.62) |
| | 24 | 0.74 (0.67-0.81) | 0.58 (0.53-0.65) | 0.57 (0.53-0.63) | 0.64 (0.56-0.71) | 0.57 (0.53-0.63) | 0.58 (0.54-0.63) |
| | 48 | 0.74 (0.66-0.80) | 0.57 (0.52-0.61) | 0.63 (0.57-0.69) | 0.61 (0.55-0.67) | 0.57 (0.52-0.62) | 0.64 (0.60-0.70) |
| | 62 | 0.73 (0.66-0.79) | 0.57 (0.53-0.62) | 0.65 (0.60-0.71) | 0.60 (0.54-0.66) | 0.57 (0.53-0.63) | 0.65 (0.60-0.71) |
| 50 | 12 | 0.78 (0.74-0.83) | 0.67 (0.62-0.72) | 0.56 (0.52-0.61) | 0.70 (0.64-0.74) | 0.66 (0.61-0.71) | 0.56 (0.52-0.60) |
| | 24 | 0.84 (0.79-0.88) | 0.71 (0.65-0.77) | 0.57 (0.53-0.61) | 0.74 (0.69-0.79) | 0.70 (0.65-0.75) | 0.56 (0.52-0.60) |
| | 48 | 0.82 (0.77-0.86) | 0.62 (0.57-0.68) | 0.55 (0.52-0.58) | 0.67 (0.62-0.72) | 0.60 (0.55-0.65) | 0.55 (0.51-0.59) |
| | 62 | 0.82 (0.78-0.87) | 0.60 (0.55-0.65) | 0.55 (0.52-0.59) | 0.65 (0.60-0.70) | 0.57 (0.53-0.62) | 0.56 (0.52-0.60) |
| 100 | 12 | 0.85 (0.82-0.88) | 0.77 (0.73-0.80) | 0.66 (0.63-0.70) | 0.78 (0.75-0.81) | 0.77 (0.73-0.80) | 0.66 (0.62-0.69) |
| | 24 | 0.91 (0.89-0.93) | 0.84 (0.81-0.86) | 0.69 (0.66-0.73) | 0.84 (0.81-0.87) | 0.83 (0.80-0.86) | 0.69 (0.65-0.72) |
| | 48 | 0.90 (0.87-0.93) | 0.75 (0.71-0.79) | 0.60 (0.56-0.63) | 0.77 (0.74-0.80) | 0.74 (0.70-0.78) | 0.59 (0.55-0.62) |
| | 62 | 0.90 (0.88-0.93) | 0.72 (0.68-0.76) | 0.59 (0.56-0.62) | 0.75 (0.71-0.78) | 0.71 (0.67-0.75) | 0.58 (0.54-0.61) |
| 250 | 12 | 0.90 (0.89-0.92) | 0.85 (0.84-0.87) | 0.78 (0.76-0.80) | 0.85 (0.84-0.87) | 0.85 (0.84-0.87) | 0.77 (0.76-0.79) |
| | 24 | 0.96 (0.95-0.97) | 0.92 (0.91-0.93) | 0.83 (0.82-0.85) | 0.92 (0.91-0.93) | 0.92 (0.91-0.93) | 0.83 (0.81-0.85) |
| | 48 | 0.96 (0.95-0.97) | 0.87 (0.85-0.88) | 0.76 (0.73-0.78) | 0.87 (0.85-0.89) | 0.87 (0.85-0.88) | 0.75 (0.73-0.77) |
| | 62 | 0.96 (0.95-0.97) | 0.86 (0.84-0.87) | 0.76 (0.73-0.78) | 0.86 (0.84-0.88) | 0.85 (0.83-0.87) | 0.75 (0.73-0.78) |

Supplementary Table 2: **AUC results from MVPA simulations for detection of site across multiple sample sizes (n) and number of features (p) where the predictor has effects on covariance.** For each of 1000 simulations, the data is randomly split into 50% training and 50% validation then a random forests algorithm is trained using the training set to predict either Site 1 or the presence of the binary predictor. The median AUC across these simulations are reported with lower and upper quartiles displayed in parentheses. Scenarios where CovBat outperforms ComBat are colored in blue.

| | | Site | | | Diagnosis Status | | |
|---|---|---|---|---|---|---|---|
| $n_i$ | $p$ | Unharmonized (%) | ComBat (%) | CovBat (%) | Unharmonized (%) | ComBat (%) | CovBat (%) |
| 25 | 12 | 99 | 41 | 16 | 98 | 99 | 99 |
| | 24 | 100 | 83 | 31 | 100 | 100 | 100 |
| | 48 | 100 | 76 | 8 | 99 | 99 | 99 |
| | 62 | 100 | 67 | 6 | 62 | 63 | 65 |
| 50 | 12 | 100 | 63 | 36 | 100 | 100 | 100 |
| | 24 | 100 | 95 | 61 | 100 | 100 | 100 |
| | 48 | 100 | 97 | 36 | 100 | 100 | 100 |
| | 62 | 100 | 99 | 40 | 100 | 100 | 100 |
| 100 | 12 | 100 | 63 | 40 | 100 | 100 | 100 |
| | 24 | 100 | 94 | 73 | 100 | 100 | 100 |
| | 48 | 100 | 98 | 62 | 100 | 100 | 100 |
| | 62 | 100 | 100 | 72 | 100 | 100 | 100 |
| 250 | 12 | 100 | 31 | 21 | 100 | 100 | 100 |
| | 24 | 100 | 70 | 48 | 100 | 100 | 100 |
| | 48 | 100 | 70 | 33 | 100 | 100 | 100 |
| | 62 | 100 | 88 | 50 | 100 | 100 | 100 |

Supplementary Table 3: **MANOVA rejection rates for associations with site and diagnosis status across multiple sample sizes (n) and number of features (p) in the Predictor Affects Mean simulations.** For each of 1000 simulations, MANOVA is performed separately for site and diagnosis using Pillai's trace. The rejection rate across these simulations is reported as the percentage of $p$-values less than 0.05.

| | | Site | | | Diagnosis Status | | |
|---|---|---|---|---|---|---|---|
| $n_i$ | $p$ | Unharmonized (%) | ComBat (%) | CovBat (%) | Unharmonized (%) | ComBat (%) | CovBat (%) |
| 25 | 12 | 96 | 10 | 1 | 92 | 94 | 96 |
| | 24 | 100 | 44 | 4 | 100 | 100 | 100 |
| | 48 | 100 | 42 | 1 | 96 | 97 | 98 |
| | 62 | 100 | 40 | 1 | 62 | 64 | 66 |
| 50 | 12 | 100 | 14 | 2 | 100 | 100 | 100 |
| | 24 | 100 | 64 | 10 | 100 | 100 | 100 |
| | 48 | 100 | 73 | 3 | 100 | 100 | 100 |
| | 62 | 100 | 86 | 3 | 100 | 100 | 100 |
| 100 | 12 | 100 | 12 | 2 | 100 | 100 | 100 |
| | 24 | 100 | 60 | 11 | 100 | 100 | 100 |
| | 48 | 100 | 68 | 5 | 100 | 100 | 100 |
| | 62 | 100 | 87 | 9 | 100 | 100 | 100 |
| 250 | 12 | 100 | 2 | 0 | 100 | 100 | 100 |
| | 24 | 100 | 11 | 2 | 100 | 100 | 100 |
| | 48 | 100 | 8 | 0 | 100 | 100 | 100 |
| | 62 | 100 | 18 | 1 | 100 | 100 | 100 |

Supplementary Table 4: **MANOVA rejection rates for associations with site and diagnosis status across multiple sample sizes (n) and number of features (p) in the Predictor Affects Covariance simulations.** For each of 1000 simulations, MANOVA is performed separately for site and diagnosis using Pillai's trace. The rejection rate across these simulations is reported as the percentage of $p$-values less than 0.05. Scenarios where the rejection rate is less than 0.05 are colored in blue.

| | | Site | | | Diagnosis Status | | |
|---|---|---|---|---|---|---|---|
| n | p | Unharmonized (%) | ComBat (%) | CovBat (%) | Unharmonized (%) | ComBat (%) | CovBat (%) |
| 25 | 12 | 94 | 1 | 0 | 74 | 78 | 82 |
| | 24 | 100 | 25 | 0 | 97 | 98 | 98 |
| | 36 | 100 | 48 | 1 | 99 | 99 | 99 |
| | 48 | 100 | 50 | 0 | 100 | 100 | 100 |
| | 62 | 100 | 47 | 1 | 96 | 97 | 98 |
| 50 | 12 | 100 | 1 | 0 | 99 | 99 | 99 |
| | 24 | 100 | 53 | 3 | 100 | 100 | 100 |
| | 36 | 100 | 83 | 6 | 100 | 100 | 100 |
| | 48 | 100 | 92 | 9 | 100 | 100 | 100 |
| | 62 | 100 | 99 | 16 | 100 | 100 | 100 |
| 100 | 12 | 100 | 2 | 0 | 100 | 100 | 100 |
| | 24 | 100 | 52 | 7 | 100 | 100 | 100 |
| | 36 | 100 | 86 | 18 | 100 | 100 | 100 |
| | 48 | 100 | 95 | 27 | 100 | 100 | 100 |
| | 62 | 100 | 100 | 61 | 100 | 100 | 100 |
| 250 | 12 | 100 | 0 | 0 | 100 | 100 | 100 |
| | 24 | 100 | 9 | 1 | 100 | 100 | 100 |
| | 36 | 100 | 29 | 6 | 100 | 100 | 100 |
| | 48 | 100 | 48 | 8 | 100 | 100 | 100 |
| | 62 | 100 | 83 | 29 | 100 | 100 | 100 |
| 500 | 12 | 100 | 0 | 0 | 100 | 100 | 100 |
| | 24 | 100 | 0 | 0 | 100 | 100 | 100 |
| | 36 | 100 | 0 | 0 | 100 | 100 | 100 |
| | 48 | 100 | 0 | 0 | 100 | 100 | 100 |
| | 62 | 100 | 4 | 0 | 100 | 100 | 100 |

Supplementary Table 5: **MANOVA rejection rates for associations with site and diagnosis status across multiple sample sizes (n) and number of features (p) in the Simple Covariance Effect simulations.** For each of 1000 simulations, MANOVA is performed separately for site and diagnosis using Pillai's trace. The rejection rate across these simulations is reported as the percentage of $p$-values less than 0.05. Scenarios where the rejection rate is less than 0.05 are colored in blue.
